# The genomic diversification of clonally propagated grapevines

**DOI:** 10.1101/585869

**Authors:** Amanda M. Vondras, Andrea Minio, Barbara Blanco-Ulate, Rosa Figueroa-Balderas, Michael A. Penn, Yongfeng Zhou, Danelle Seymour, Ye Zhou, Dingren Liang, Lucero K. Espinoza, Michael M. Anderson, M. Andrew Walker, Brandon Gaut, Dario Cantu

## Abstract

Vegetatively propagated clones accumulate somatic mutations. The purpose of this study was to better understand the consequences of clonal propagation and involved defining the nature of somatic mutations throughout the genome. Fifteen Zinfandel winegrape clone genomes were sequenced and compared to one another using a highly contiguous genome reference produced from one of the clones, Zinfandel 03.

Though most heterozygous variants were shared, somatic mutations accumulated in individual and subsets of clones. Overall, heterozygous mutations were most frequent in intergenic space and more frequent in introns than exons. A significantly larger percentage of CpG, CHG, and CHH sites in repetitive intergenic space experienced transition mutations than genic and non-repetitive intergenic spaces, likely because of higher levels of methylation in the region and the increased likelihood of methylated cytosines to spontaneously deaminate. Of the minority of mutations that occurred in exons, larger proportions of these were putatively deleterious when they occurred in relatively few clones.

These data support three major conclusions. First, repetitive intergenic space is a major driver of clone genome diversification. Second, clonal propagation is associated with the accumulation of putatively deleterious mutations. Third, the data suggest selection against deleterious variants in coding regions such that mutations are less frequent in coding than noncoding regions of the genome.

## Introduction

Cultivated grapevines are clonally propagated. As a result, the genome of each cultivar is preserved, except for the accumulation of mutations that accumulate over time and can generate distinguishable clones [1-4]. Somatic mutations are responsible for several notable phenotypes. For example, a single, semi-dominant nucleotide polymorphism can affect hormone response [5] and recessive insertion of the *Gret1* retrotransposon in the promoter of the *VvmybA1* transcription factor inhibits anthocyanin accumulation in white varieties [6], as do additional mutations affecting the color locus [7-10]. The fleshless fruit of an Ugni Blanc clone and the reiterated reproductive meristems observed in a clone of Carignan are both caused by dominant transposon insertion mutations [11,12]. In citrus, undesirable mutations can be unknowingly propagated that render fruit highly acidic and inedible [13,14]. Interestingly, somatic mutations in plum are associated with a switch from climacteric to non-climacteric ripening behavior [15].

There is limited understanding and evidence of the extent, nature, and implications of the somatic mutations that accumulate in clonally propagated crops [16]. Genotyping approaches based on whole genome sequencing make it possible to identify genetic differences without predefined markers [17-19] and expedite learning the genetic basis of valuable traits and developmental processes [15,20]. Still, few previous studies have used genomic approaches to study somatic variations among clones [17-21]. The first to publish a genome-wide exploration of somatic variation in grapevine was Carrier *et al.* (2012), finding that transposable elements were the largest proportion of somatic mutation types affecting four Pinot Noir clones [18].

Whole genome sequencing was also used to study structural variations and complex chromosomal rearrangements in Tempranillo, comparing diverse accessions of phenotypically distinct Tempranillo Tinto and Tempranillo Blanco to better understand the basis of somatic mutations giving rise to red versus white fruit [20]. Genomic tools could be used to comprehensively describe the extent of somatic mutations and infer the processes affecting clone genomes.

Mutations occur in somatic cells that proliferate by mitosis. These can occur by a variety of means, including single base-pair mutations [22,23] that are more prevalent in repetitive regions because methylated cytosines passively deaminate to thymines [24-26], polymerase slippage that drives variable microsatellite insertions and deletions [27], and larger structural rearrangements and hemizygous deletions [10,20]. Transposable elements are also a major source of somatic mutations in grapevines [18], though transcriptional and post-transcriptional mechanisms exist to prevent transposition and maintain genome stability [28-31]. Notably, methylation of transposable elements is one specific mechanism that prevents transposition, which establishes a tradeoff, then, between methylation and the transposition of mobile elements.

At the cellular level, distinct clones can emerge following a mutation in a shoot apical meristem that spreads throughout a single cell layer, creating periclinal chimeras. This chimera is stable for Pinot Meunier, a clone of Pinot Noir with distinct L1 and L2 layers in shoots [3]. Each cell layer in a stratified apical meristem like that observed in grape [32] is developmentally distinct. The distinct cell layers will remain so provided cell divisions occur anticlinally. But, periclinal divisions and cellular rearrangements can result in the homogenization of a mutant genotype across cell layers [33]. This is the case for green-yellow bud sports of the grey-fruited Pinot Gris, wherein sub-epidermal white cells invaded and displaced epidermal pigmented cells [9]. In contrast to replacement (L1 cells invade L2), displacement is likely more common because of the relative disorganization of the inner cell layers [32,33].

Meristem architecture is related to the fate of somatic mutations, as it influences the impact of these mutations and the likelihood of competition between cell lineages, also known as diplontic selection [34-36]. Provided each cellular layer is maintained by anticlinal divisions, deleterious mutations can be preserved in periclinal chimeras [35,37]. In addition, the predominance of “hidden”, heterozygous recessive somatic mutations [2,37] may further shield somatic mutations from selective forces. These factors are permissive of the accumulation of somatic mutations. Diplontic selection could occur if periclinal cell divisions result in the invasion of one cell layer by cells from another [34,35]. This mechanism could oppose the accrual of deleterious mutations expected by Muller [38,39]. A recent study of the long-lived pedunculate oak described substantial intra-organismal genetic variation, but did not draw conclusions about the contribution of somatic variations to large-scale oak evolution [21]. Evidence of diplontic selection in plants is remarkably scarce [37], though its likelihood given different circumstances has been modeled [34,35,40]. Given the prevalence of chimerism and rearrangements documented in the model [9,33], grapevine is a suitable model for investigating the possibility of selection during vegetative propagation.

Zinfandel is the third-most cultivated wine grape in California [41,42] DNA profiling produced evidence that Zinfandel is synonymous with Primitivo grown in Italy [43] and Croatian Pribidrag and Crljenak Kastelanski [44]. Historical records plus the cultivation of closely related cultivars support Croatia as the likely origin of Zinfandel [44-47] and also that Primitivo was likely brought to the Gioia del Colle region in Italy by Benedictine monks in the 17^th^ century [3,48]. The reported variability in Zinfandel [49-51], including subtle variability in phenolic metabolites (Additional file 1), and its long history of cultivation make it a useful model for studying clonal variation in grapevine, specifically, and the nature of the accumulation of somatic mutations in clonally propagated crops, generally.

The purpose of this study was to better understand the nature of the somatic variations that occur during clonal propagation. Representatives from at least a portion of Zinfandel’s history [44-47] from Croatia, Italy, and California were sequenced and compared using Zin03 as reference. First, we show that intergenic space drives clonal diversification. As previously reported, transposable element insertions varied among clones [18]. This report expands that understanding to implicate methylation as an indirect driver of clonal diversification; rare somatic heterozygous SNPs were most observed in the repetitive intergenic regions, likely because of the high levels of transposition-inhibiting methylation and associated transition mutations that are prevalent there. Second, the data support an important component of Muller’s ratchet [38], that asexually propagated organisms accumulate deleterious mutations. Third, somatic mutations were relatively scarce in the coding regions of genes relative to introns and intergenic space, suggesting some degree of negative selection against deleterious mutations.

## Results

### Zinfandel genome assembly, annotation, and differences between haplotypes

The clone used for the genome assembly, Zinfandel 03 (Zin03), was acquired by FPS in 1964 from the Reutz Vineyard near Livermore, California that was planted during Prohibition (1920 – 1933) [52]. Zin03 was sequenced using Single Molecule Real-Time (SMRT; Pacific Biosciences) technology at ∼98x coverage and assembled using FALCON-unzip [53], a diploid-aware assembly pipeline. The genome was assembled into 1,509 primary contigs (N50 = 1.1 Mbp) for a total assembly size of 591 Mbp, similar to the genome size of Cabernet Sauvignon (590 Mbp) [53] and larger than Chardonnay (490Mb) [19] and PN40024 (487 Mb) [54]. Fifty two percent of the genome was phased into 2,246 additional phased sequences (haplotigs) where the homologous chromosomes were distinguishable with an N50 of ∼442 kbp (Table 2). A total of 53,560 complete protein-coding genes were annotated on the primary (33,523 genes) and haplotig (20,037 genes) assemblies (Table 2).

**Table 1.**
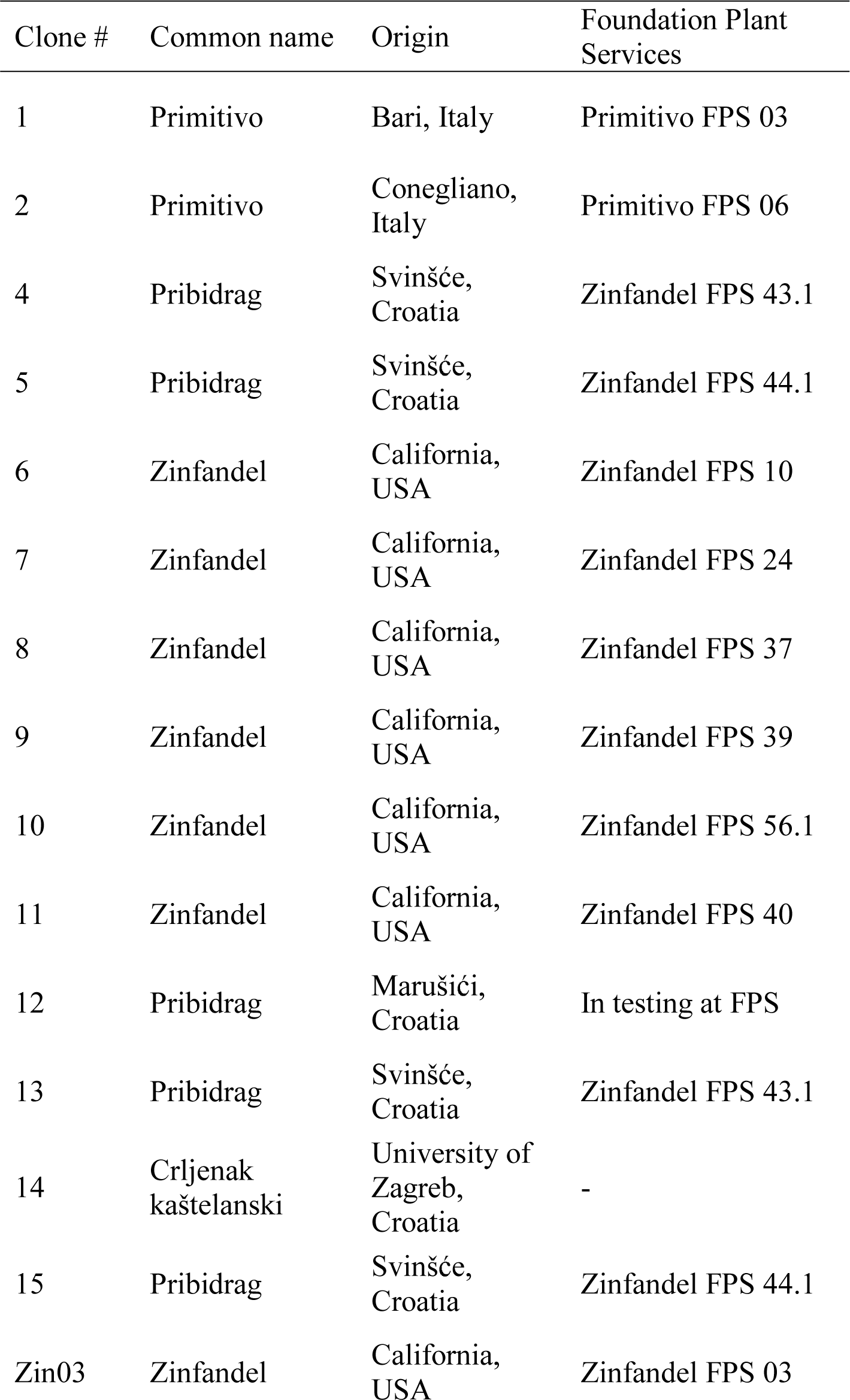
Clone identifying information

| Clone # | Common name | Origin | Foundation Plant Services |
| --- | --- | --- | --- |
| 1 | Primitivo | Bari, Italy | Primitivo FPS 03 |
| 2 | Primitivo | Conegliano, Italy | Primitivo FPS 06 |
| 4 | Pribidrag | Svinšće, Croatia | Zinfandel FPS 43.1 |
| 5 | Pribidrag | Svinšće, Croatia | Zinfandel FPS 44.1 |
| 6 | Zinfandel | California, USA | Zinfandel FPS 10 |
| 7 | Zinfandel | California, USA | Zinfandel FPS 24 |
| 8 | Zinfandel | California, USA | Zinfandel FPS 37 |
| 9 | Zinfandel | California, USA | Zinfandel FPS 39 |
| 10 | Zinfandel | California, USA | Zinfandel FPS 56.1 |
| 11 | Zinfandel | California, USA | Zinfandel FPS 40 |
| 12 | Pribidrag | Marušići, Croatia | In testing at FPS |
| 13 | Pribidrag | Svinšće, Croatia | Zinfandel FPS 43.1 |
| 14 | Crljenak kaštelanski | University of Zagreb, Croatia | - |
| 15 | Pribidrag | Svinšće, Croatia | Zinfandel FPS 44.1 |
| Zin03 | Zinfandel | California, USA | Zinfandel FPS 03 |

**Table 2.** Summary statistics of the Zinfandel genome assembly and annotation.

|  | Primary | Haplotig |
| --- | --- | --- |
| Total length | 591,171,721 | 306,029,957 |
| Number of contigs | 1,509 | 2,246 |
| N50 | 1,062,797 | 442,393 |
| N75 | 366,308 | 185,785 |
| L50 | 154 | 200 |
| L75 | 395 | 463 |
| Median contig length (bp) | 161,249 | 37,307 |
| Longest contig (bp) | 7,901,503 | 2,609,171 |
| Shortest contig (bp) | 17,787 | 1,970 |
| Average GC content (%) | 34.45% | 34.37% |
| Number of genes | 33,523 | 20,037 |
|  | *Total* | *Average per gene* |
| Number of exons | 244,880 | 4.57 |
| Number of introns | 191,320 | 3.57 |
|  | *Average (bp)* | *Maximum (bp)* |
| mRNA lengths | 4,166 | 94,143 |
| Exon lengths | 245.79 | 7,992 |
| Intron lengths | 191,320 | 41,647 |
| Intergenic distances | 10,309 | 302,473 |

Of the 20,037 genes annotated on the haplotig assembly, 18,878 aligned to the primary assembly, leaving 1,159 genes that may exist hemizygously in the genome due to structural variation between homologous chromosomes or because of substantial divergence in sequence between haplotypes. These genes were annotated with a broad variety of putative functions, including biosynthetic processes, secondary metabolism, and stress responses. Long reads were mapped to both the primary and haplotig assemblies to evaluate the circumstances that explain the differences between haplotypes. Structural variants (SVs) between the haplotypes were examined by mapping long SMRT sequencing reads onto Zin03’s primary and haplotig assemblies with NGMLR and calling SVs with Sniffles [55].

A total of 22,399 SVs accounted for 6.94% (41.0 / 591 Mbp) of the primary assembly’s length and 6.02% (8.4 / 139 Mbp) of the primary assembly’s gene-associated length (Fig. 1a, Table 3). SVs intersected 4,559 genes in the primary assembly (13.6% of primary assembly genes) and 390 SVs spanned more than one gene. Manual inspection of the long reads aligned to the primary assembly support that large, heterozygous deletions and inversions occurred in the Zin03 genome that were either inherited from different structurally distinct parents or arose during clonal propagation (Fig. 1b,c,d). Importantly, there was substantial hemizygosity in the genome, with long reads supporting deletions affecting 2,521 genes and 4.56% of the primary assembly’s length (Table 3).

**Table 3.** Sniffles analysis of structural variation between cultivars and between Zinfandel parental haplotypes

|  | Cabernet Sauvignon vs. Zinfandel |  |  |  |  | Zinfandel haplotig vs. Zinfandel primary |  |  |  |  |
| --- | --- | --- | --- | --- | --- | --- | --- | --- | --- | --- |
|  | *Median Size (bp)* | *Count* | *Genes* | *Total SV size (Mb)* | *% genome* | *Median Size (bp)* | *Count* | *Genes* | *Total SV size (Mb)* | *% genom* |
| Deletions | 196 | 46,363 | 9,219 | 115.0 | 12.82 | 203 | 12,031 | 2,521 | 26,953,558 | 4.56 |
| Duplications | 5,518 | 2,884 | 3,286 | 48.7 | 5.43 | 1,966 | 553 | 535 | 7,604,041 | 1.29 |
| Insertions | 88 | 37,407 | 5,225 | 23.9 | 2.66 | 92 | 9,647 | 2,081 | 5,594,259 | 0.95 |
| Inversions | 6,037 | 607 | 1,440 | 20.6 | 2.30 | 3,592 | 111 | 391 | 5,521,214 | 0.93 |
| Duplicated Insertions | 339 | 9 | 2 | 0.0439 | 0.0049 | 385 | 3 | 2 | 6,861 | 0.0012 |
| Inverted Duplications | 293 | 65 | 12 | 0.0418 | 0.0047 | 113 | 54 | 11 | 12,930 | 0.0022 |

**Figure 1.**
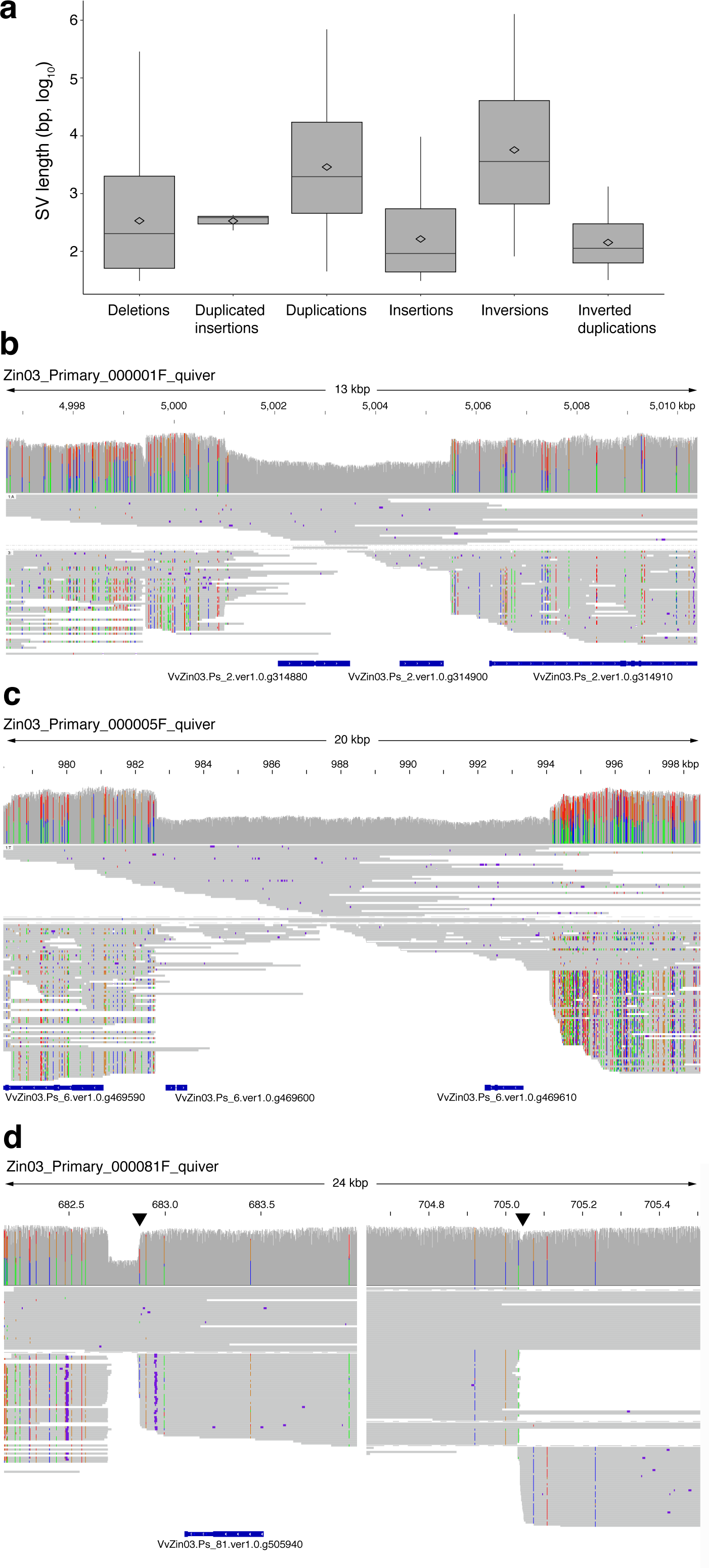
Structural variation between Zin03 haplotypes. **a**. Distribution of structural variation sizes. Boxplots show the 25^th^ quartile, median, and 75^th^ quartile for each type of SV. Whiskers are 1.5^Inter-Quartile Range^. Diamonds indicate the mean log_10_(length) of each type of SV; **b**,**c**,**d.** Examples of heterozygous structural variants between haplotypes that intersect genes. For each reported structural variation, (from top to bottom) the coverage, haplotype-resolved alignment of reads, and the genes annotated in the region are shown; **b.** 4 kbp heterozygous deletion of two genes; **c.** 11 kbp heterozygous deletion of two genes; **d.** 22 kbp inversion that intersects a single gene. Triangles indicate boundaries of the inversion. A gap is shown rather than the center of the inverted region.

Next, we considered whether specific structural variation could account for the 1,159 genes uniquely found in the haplotig assembly. Three hundred eighty-two genes of the previously mentioned 1,159 genes that uniquely exist within the haplotig assembly intersected structural variations. Two hundred ninety of these intersected deletions, accounting for the failure to identify them on the primary assembly. Some of the haplotig genes that failed to map to the primary assembly intersected additional types of SVs, including duplications (80 genes), insertions (89 genes), and inversions (16 genes).

These results reveal structural differences between Zinfandel’s haplotypes. These differences could have been inherited and/or could have occurred during clonal propagation. Overall, these structural variations affected 4,559 primary assembly genes. Importantly, these data show that a notable portion of the primary assembly’s length (4.56%) is hemizygous.

### Differences in structure and gene content between Zinfandel and other grape genomes

The Zin03 genome was compared to PN40024 and Cabernet Sauvignon to identify cultivar-specific genes that may contribute to Zinfandel’s characteristics. PN40024 is the inbred line derived from Pinot Noir used to develop the first grape genome reference [54] and Cabernet Sauvignon (CS08) was recently used to construct the first diploid, haplotype-resolved grape genome for which long reads are available [53]. Overall, 1,801 genes were not shared between all three genotypes (Zin03, Pinot Noir, and Cabernet Sauvignon; Fig. 2a). Three hundred nine protein coding genes were found uniquely in Zin03 relative to PN40024 and CS08; 223 were annotated on the primary assembly and 86 were annotated on the haplotigs (Fig. 2a, Additional file 2). These genes had a panoply of functions that included but were not limited to nucleotide binding (60 genes), protein binding (58 genes), stress response (34 genes), and kinases (28), and were associated with membranes (48 genes), signal transduction (23 genes), carbohydrate metabolism (12 genes), and lipid metabolism (8 genes; Additional file 2).

**Figure 2.**
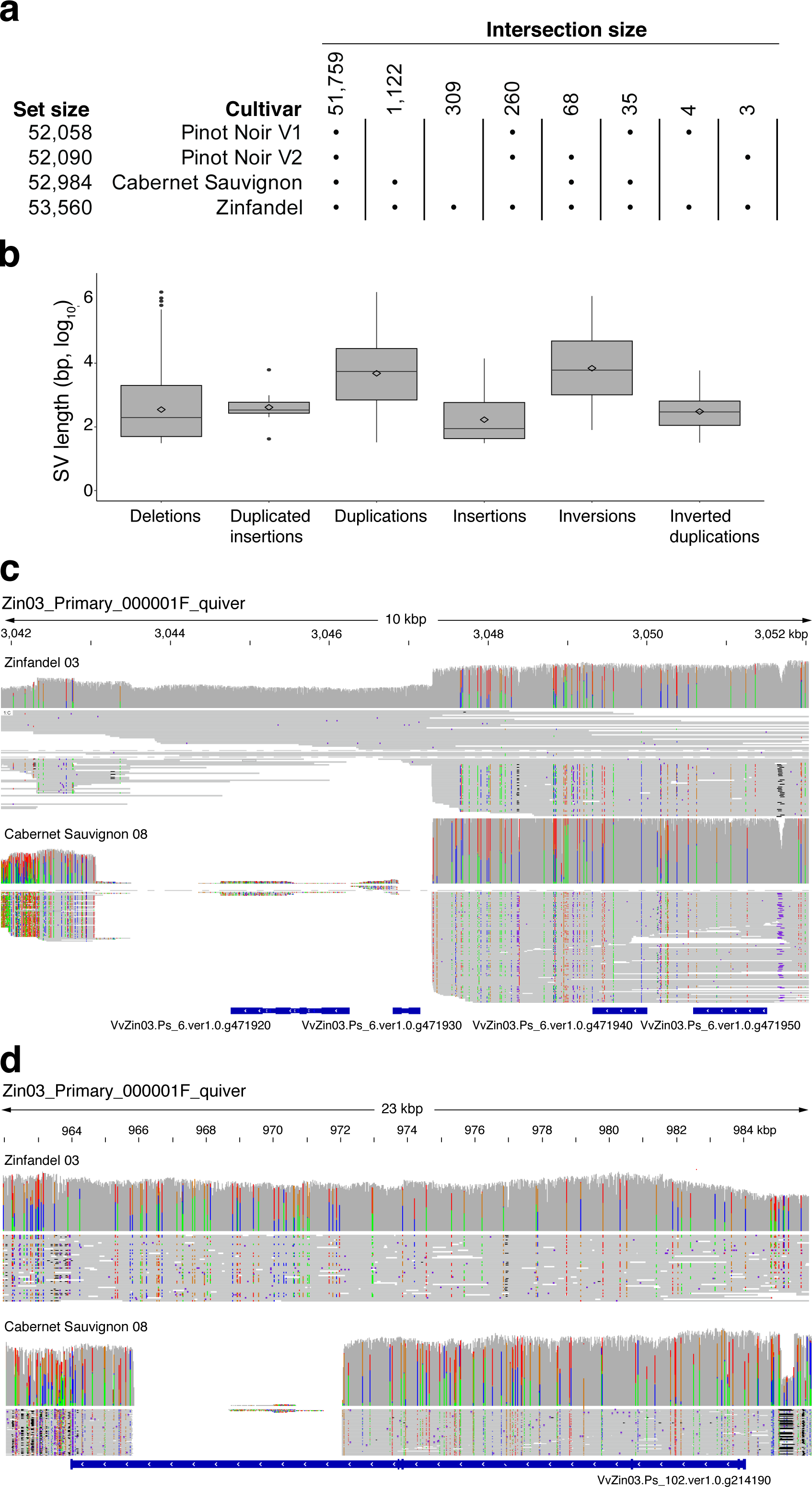
Gene content and structural variability between Zin03 and other *V. vinifera* genomes. **a.** Uniquely occurring Zinfandel genes and the number of Zinfandel genes that align well to other cultivars with >=80% identity and reciprocal coverage. The total number of hits (or total gene content for Zin03) is indicated by the “Set Size” and the exclusive hits for each intersection is indicated as the “Intersection Size”; **b. Boxplot shows the sizes of structural variations; c**,**d.** Selected deletions in Cabernet sauvignon relative to Zin03 that intersect genes. For each reported deletion, (from top to bottom) the coverage of reads over the region by long Zinfandel and Cabernet Sauvignon reads, haplotype-resolved alignment of the reads, and the genes annotated in the region are shown; **b.** Two genes are completely deleted in Cabernet Sauvignon relative to Zinfandel and are deleted in one Zinfandel haplotype; **c.** One gene contains a homozygous partial deletion in Cabernet Sauvignon.

Structural differences between Zin03 and CS08 were explored in more detail by mapping the long SMRT reads of CS08 onto Zin03’s primary and haplotig assemblies with NGMLR and calling SVs with Sniffles (Fig. 2b, Table 3). Overall, these SVs corresponded to 17.74% (159/ 897 Mbp) of the Zin03 assembly’s total length, 12.5% of its total protein-coding regions (28 / 223 Mbp), and 25.6% of all Zin03 genes. SVs affected 9,885 genes in the primary assembly and 3,804 genes in the haplotigs. Manual inspection of the alignment of long CS08 reads to Zin03’s primary assembly support that large SVs exist between the two genotypes (Fig. 2c,d). Next, we considered whether specific structural variation called by Sniffles could account for the 576 Zin03 genes absent from CS08 according to the reciprocal mapping analysis (Fig. 2a). Of these 576 Zinfandel genes, 268 genes intersected 454 deletions supported by long CS08 reads aligned to Zin03.

Though Zinfandel had few unique genes, high levels of structural variation between Zinfandel (Zin03) and Cabernet Sauvignon (CS08) were observed and these affected considerable protein-coding regions of the genome. These results justify constructing a Zinfandel-specific reference to better capture genomic variability among Zinfandel clones that could otherwise be missed, particularly if an alternative reference lacks sequences present in Zinfandel.

### Relatedness among Zinfandel clones

Fifteen Zinfandel clones, including Zin03, were sequenced using Illumina. The resulting reads were aligned to the Zin03 primary assembly to characterize SNPs, small INDELs, variable transposon insertions, and large structural variants. Principal Component Analysis (PCA) of variants among the clones showed no clear pattern in their relationships to one another based on their recorded origins prior to acquisition (Fig. 3a). The ambiguity surrounding the travels and histories of these clones means that it should not be taken for granted that the Californian selections, for example, ought to be more closely related to one another than to the Italian or Croatian selections. Notably, Pribidrags 5 and 15, which have a known and close relationship, do not co-localize in the PCA (Fig. 3a, Table 1).

**Figure 3.**
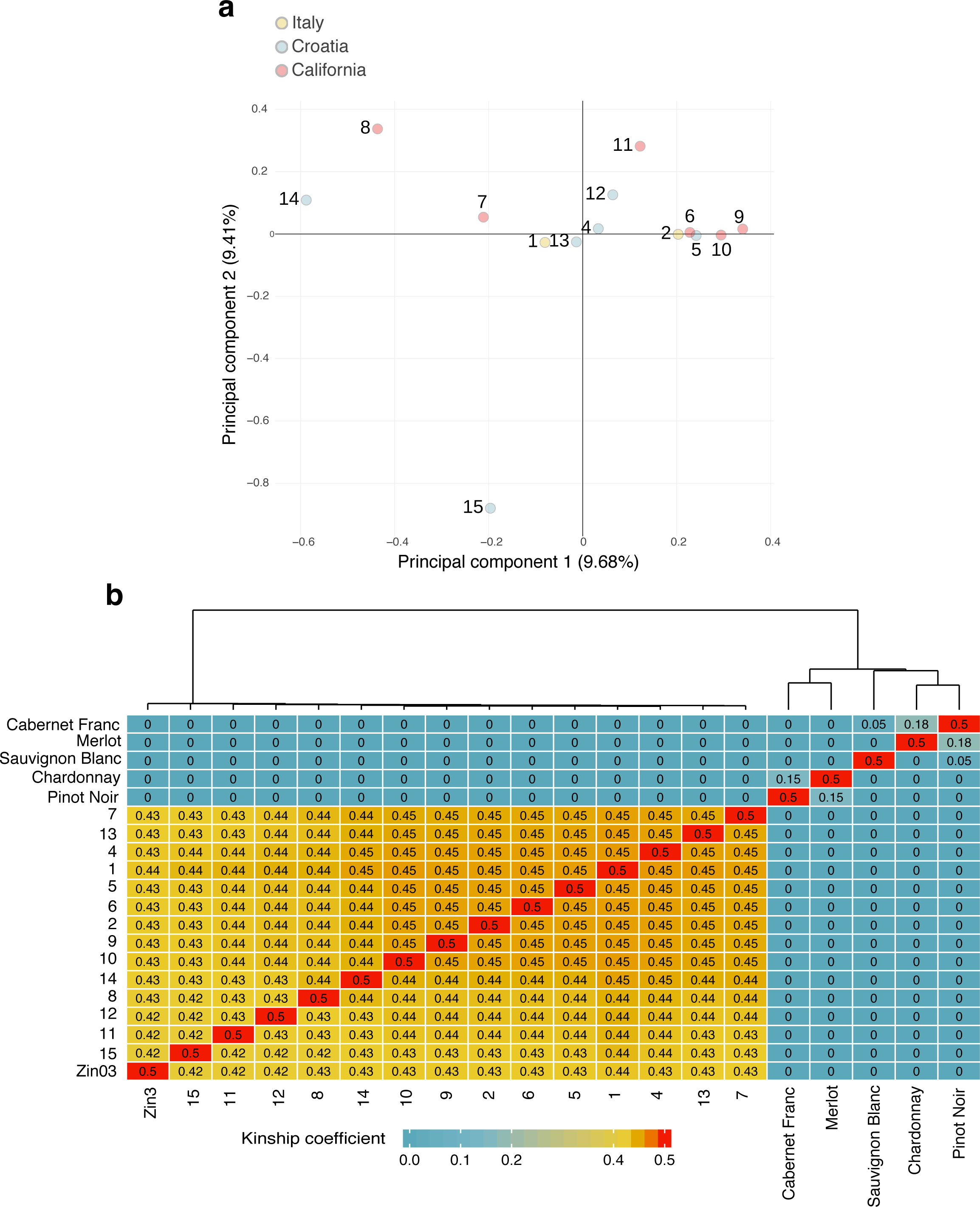
The relationships between Zinfandel selections. **a.** Principal component analysis of Zinfandel selections based on SNP data. Zin03 was not included in the analysis; **b.** Kinship analysis of Zinfandel selections and other cultivars with known relationships based on SNP data and outside of annotated repeats. The Kinship coefficient, PHI, is shown, as well as a dendrogram constructed by hierarchically clustering genotypes using their kinship coefficients.

A kinship analysis [56] was then used to quantitatively assess the relationships between the Zinfandel selections. These values range from zero (unrelated) to 0.5 (self). Additional cultivars were included in the analysis with known relationships to help contextualize the differences between clones and the integrity of the analysis (Fig. 3b). Cabernet Franc and Merlot have a parent – offspring relationship, as do Pinot Noir and Chardonnay [57,58]. These pairs had kinship coefficients of 0.15 and 0.18, respectively (Fig. 3b). As a possible grandparent of Sauvignon Blanc, Pinot Noir had a kinship coefficient of 0.05 with Sauvignon blanc [59,60]. Zinfandel selections had kinship coefficients between 0.42 and 0.45; this is likely because of the accrual of somatic mutations among clones (Fig. 3b).

Across the Zinfandel clones, the median number of homozygous and heterozygous variants called relative to Zin03 were 37,437 and 718,174, respectively. Between 10-fold and 27-fold more heterozygous variants were called than homozygous variants in each clone, and less than 10% of sites did not share the Zin03 reference allele (Additional file 3).

### Clonal versus cultivar genetic variability

Overall, an average of 761,948 variant sites were identified in individual Zinfandel clones when short reads were mapped on the Zin03 primary assembly. On average, 6,153,830 variant sites were identified in other cultivars (Pinot noir, Chardonnay, Sauvignon Blanc, Merlot, Cabernet Franc) relative to Zin03 (Additional file 3). Both of these figures excluded heterozygous sites at which the diploid genotype called for a given sample was identical to that called for Zin03.

Variants were 7.9X more frequent in other cultivars relative to Zin03 than for Zinfandel clones; on average, mutations in clones occurred once every 723 bases and once every 92 bases in other cultivars (Additional file 3). However, the ratio of transitions to transversion mutations and the proportions of the severities of the predicted variant effects were similar for both groups (Additional file 3). The normalized count of variants differed between cultivars and Zinfandel clones on the basis of variants’ location in the genome, the type of variant, and the zygosity of the variant (Fig. 4).

**Figure 4.**
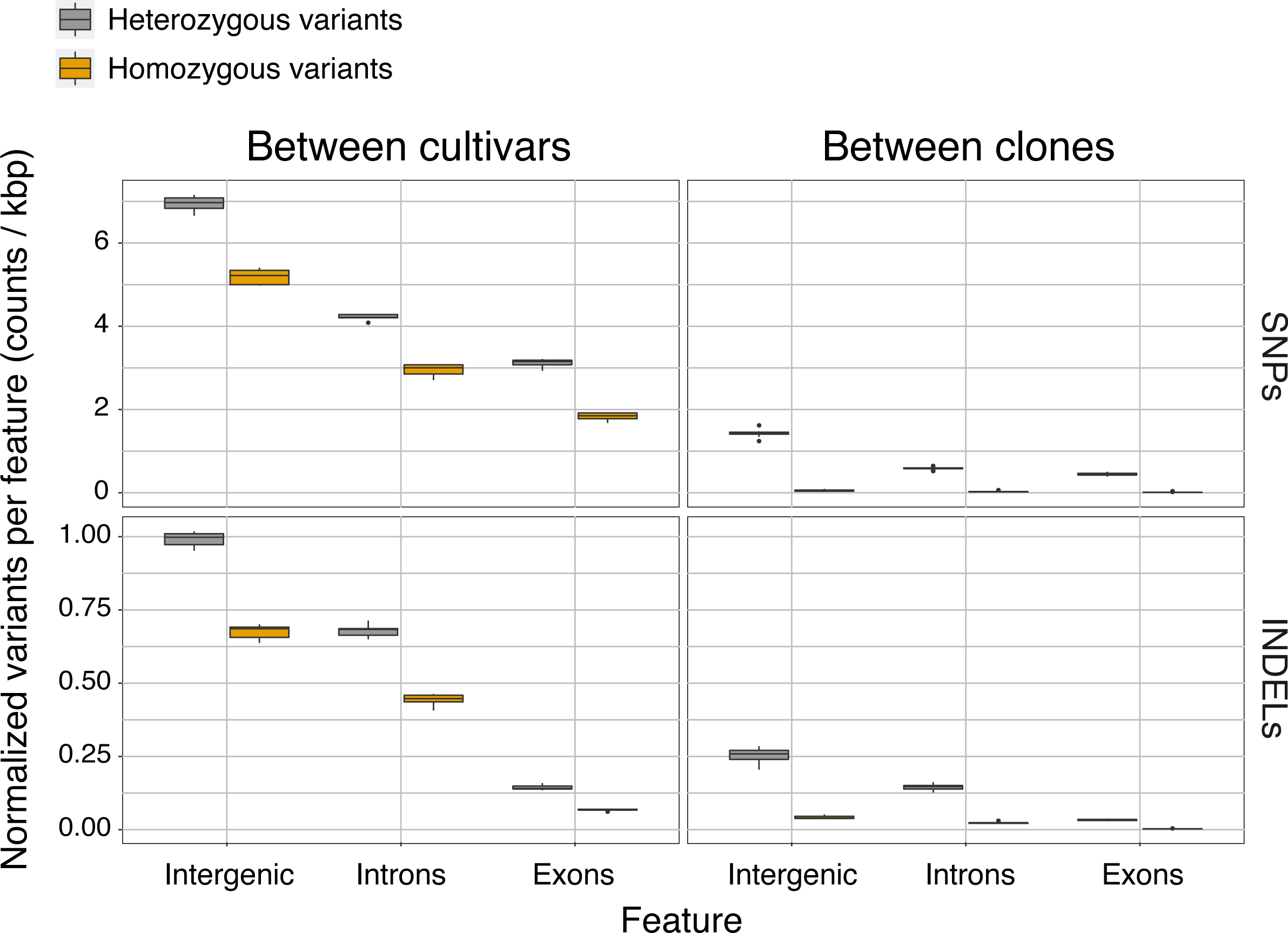
Characterization of variants and their frequency among Zinfandel selections and other *vinifera* cultivars (Pinot Noir, Chardonnay, Merlot, Cabernet Franc, and Sauvignon Blanc). The normalized rate of variants (number of variants divided by the total feature length in the genome * 1k) by type (SNP, INDEL), feature (Intergenic, Intron, Exon), and genotype (Non-Zinfandel Cultivars, Zinfandel selections). Boxplots show the 25^th^ quartile, median, and 75^th^ quartile.

Variants in non-Zinfandel cultivars and heterozygous variants among Zinfandel clones were significantly more prevalent in intergenic space than introns and exons and significantly more prevalent in introns than exons (Tukey HSD, *p <* 0.01). Unlike homozygous variants between cultivars and as expected, homozygous variants were rare among clones (Fig. 4, Additional file 3). Still, the normalized count of homozygous INDELs in intergenic space, introns, and exons were significantly different among Zinfandel clones (Tukey HSD, *p <* 0.01), as were the normalized count of homozygous intergenic versus genic (exons and introns) SNPs (Tukey HSD, *p <* 0.01). The normalized count of homozygous SNPs in exons and introns were not significantly different in Zinfandel clones (Tukey HSD, *p* > 0.01). The accrual of predominantly heterozygous and likely recessive variants [2] is consistent with what would be expected given physically separate homologous chromosomes and the absence of sexual reproduction. The differences in mutation abundances observed were initially surprising; if somatic mutations occurred randomly and absent mechanisms that make certain sites more or less susceptible to mutation, then different regions of the genome should have had equal levels of mutations. This was not the case (Figure 5).

**Figure 5.**
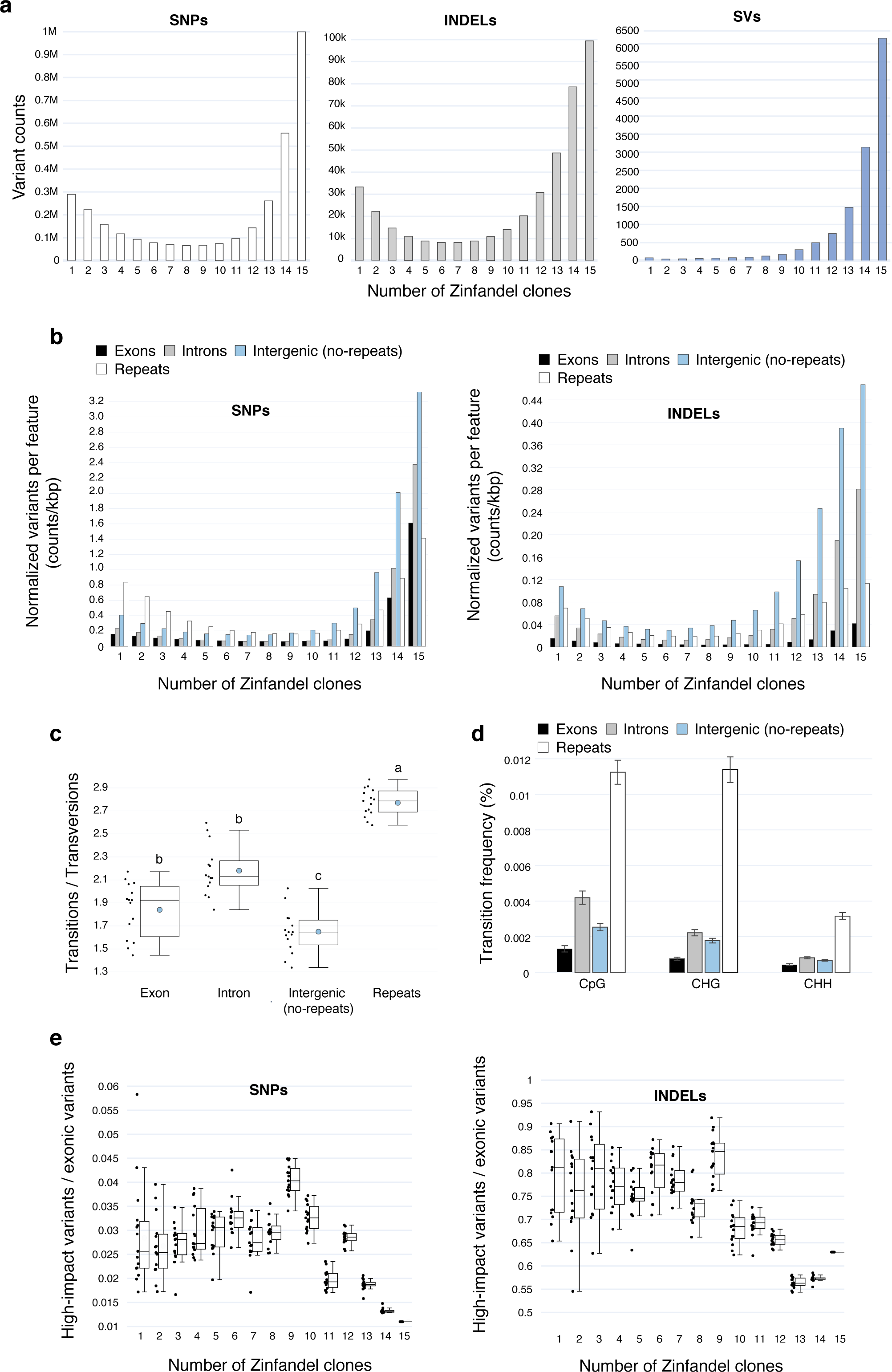
The abundance and impact of shared and unique heterozygous mutations among Zinfandel clones. **a.** The number of heterozygous SNPs, INDELs, and SVs are shared by N Zinfandel clones; **b.** The number of SNPs and INDELs shared by N clones in exons, introns, intergenic repeats (“Repeats”), and non-repetitive intergenic space; **c.** The ratio of transitions (Tr) to transversions (Tv) for heterozygous SNPs that uniquely occur in single Zinfandel clones and in different genome features. Different letters correspond to significant differences in Tr/Tv rates between features (ANOVA, Tukey HSD, *p* < 0.01); **d.** The percentage of CpG, CHG, and CHH in exons, introns, intergenic repeats (“Repeats”), and non-repetitive intergenic space that experiences transition mutations. Comparisons were made between features for each type of C-repeat separately. Different letters correspond to significant differences (Tukey HSD, *p* < 0.01); **e.** Proportion of exonic SNPs and INDELs that are deleterious and shared by N Zinfandel clones

### The accrual of somatic mutations in Zinfandel clones

Heterozygous sites found among the 15 Zinfandel clones ought to be a mixture of sites inherited from their shared ancestral plant and somatic mutations that arose during clonal propagation. To better understand the nature of somatic mutations, the data were handled slightly differently than they were to construct Figure 4; all 15 Zinfandel clones were included and all heterozygous calls were considered, even if all genotypes were identically heterozygous. Thirty percent of heterozygous SNPs, 24% of heterozygous INDELs, and 47% of heterozygous structural positions were shared by all 15 Zinfandel clones (Fig. 5a). Because all clones are identically heterozygous at these loci, these variants are those inherited from Zinfandel’s parents.

Individual and subsets of Zinfandel clones accumulated heterozygous mutations as clonal propagation occurred (Fig. 5a). Thirteen percent and 16% of heterozygous INDELs and SNPs, respectively, and 1% of large (>50 bp) structural variants occurred in only one or two clones (Fig. 5a). The distribution of SVs called by Delly is markedly different than those of SNPs and INDELs (Fig. 5a). For both SNPs and INDELs, there were 3 and 3.5-fold as many heterozygous variants shared by all 15 clones as there were uniquely occurring variants; there were 71.5-fold more structural variants shared by all clones than there were unique variants in individual clones (Fig. 5a). This might imply that the mechanisms that give rise to small mutations are more common among clones than the large-scale changes associated with SVs.

The distribution of unique and shared heterozygous INDELs in exons, introns, repetitive, and non-repetitive intergenic spaces were not equal (Fig. 5b). The distribution of INDELs in exons was significantly different than the distributions of INDELs in each other feature considered (Kolmogorov-Smirnov Test, *p* < 0.01). Similarly, the distributions SNPs in genic (exons, introns) and intergenic (repetitive, non-repetitive) regions were not equal (Fig. 5b). Shared heterozygous SNPs were most common in intergenic non-repetitive regions and introns and least common in exons and repetitive intergenic regions (Fig. 5b). Interestingly, unique heterozygous SNPs occurred at high rates in repetitive intergenic regions (Fig. 5b).

That shared heterozygous sites are mostly in non-repetitive intergenic space and unique heterozygous sites are mostly in repetitive space may have to do with the increased likelihood that methylated cytosines spontaneously deaminate and the prevalence of methylated repetitive sequences in those regions [22,25,29,30]. This is also supported by the significantly higher ratio of transitions to transversions in repetitive intergenic regions than in exons, introns, and non-repetitive intergenic space (Fig. 5c). Furthermore, the mean percentage of CpG, CHG, and CHH sites affected by transition mutations was significantly higher in repetitive intergenic space than genic and non-repetitive intergenic spaces (Fig. 5d; Tukey HSD, *p* < 0.01). The mean percentage of CpG sites affected by transition mutations was also significantly higher in introns than exons (Tukey HSD, *p* < 0.01). Compatible with this hypothesis, INDELs, which should not increase in frequency due to methylation, did not occur preferentially in repeats (Fig. 5b).

The impact of specific variants also varied with their prevalence among the clones (Fig. 5e). “High impact” mutations were predicted by SNPEff [61]. The high impact mutations identified in these data included exon losses, start and stop site gains and losses, frameshifts, gene fusions, splice acceptor mutations, and splice donor mutations. These mutations are predicted to be deleterious because of their disruptive effects on the coded protein. For these reasons, we designated such mutations as putatively deleterious in this manuscript. These were counted for each Zinfandel clone relative to Zin03. Relatively low proportions of heterozygous variants shared by all Zinfandel clones were putatively deleterious. In contrast, larger proportions of exonic SNPs and INDELs that occurred in individual or subsets of clones were putatively deleterious (Fig. 5e).

Together, these results show that mutations associated with clonal propagation are most numerous outside of coding regions of the genome, indicating that clone genomes diversify most rapidly in the intergenic space, particularly in repetitive and likely methylated regions (Fig. 5). Though a minority of somatic mutations occurred in exons, we show that exonic mutations that occur in few or individual clones are more often deleterious than exonic heterozygous variants shared by all or most clones. In other words, clonal propagation is associated with the accumulation of putatively deleterious heterozygous mutations.

### Zinfandel clones incur unique transposon insertions

Transposable element insertions (TEI) contribute to somatic variation in grape [6,11,12,18]. Relative to Zin03, 1,473 TEI were identified among the Zinfandel clones. A large fraction of TEI (26.7%) occurred uniquely in individual clones (Fig. 6a) and included 325 retrotransposons, mostly Copia and Gypsy LTRs, and 69 DNA-transposons (Fig. 6b). Because uniform loci are excluded, in-common TEI were not captured when clones were compared to Zin03. Comparing the clones relative to PN40024, however, revealed that the majority (64.8%) of TEI were shared among the 15 Zinfandel clones. Five hundred thirty TEI occurred in only one, two or three clones (Fig. 6a). This result supports the derivation of these selections from a common ancestral plant and the accumulation of somatic variations over time.

**Figure 6.**
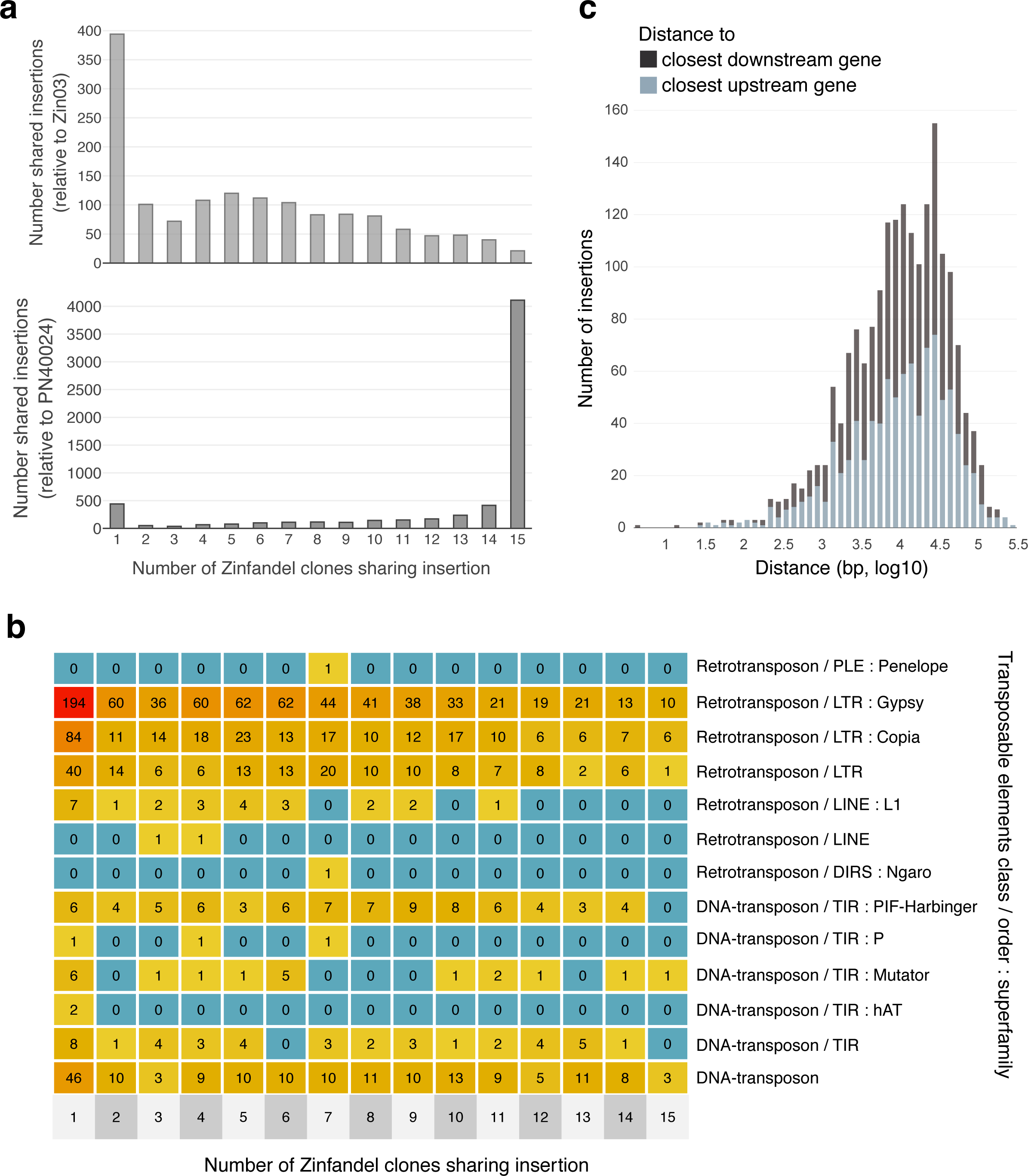
Transposable element insertions among Zinfandel selections. **a.** Transposable element insertions shared among N Zinfandel selections relative to Zin03 and PN40024; **b.** Types of transposable element insertions shared by N Zinfandel selections;**c.** The proximity of intergenic transposable element insertions to genes

In addition to being suggestive of their shared heritage, the positions of these insertions and their proximity to coding genes were notable. Three-hundred forty-seven TEI occurred within 314 coding genes. The remaining 938 TEIs were in intergenic regions (Fig. 6c). The median upstream and downstream distance of intergenic TEs from the closest feature were 11,811 and 11,279 base-pairs, respectively, and 25% of TEI were less than 4,345 bases downstream of the closest feature and/or less than 3,826 bases upstream of the closest feature (Fig. 6c).

## Discussion

Consideration of the genomic differences among Zinfandel clones revealed what is likely a complex history not easily reconstructed. Analyses of the relationships between clones did not reveal groupings of clones per their recorded countries of origin. Somatic mutations may help identify individual clones but could also blur the historical relationships between them. It is also plausible that pairs of clones from any given region are not direct cuttings of one another but of Zinfandels from another region; the clones now grown in California, for example, may have been imported on numerous independent occasions from various other regions, meaning some may indeed be more closely related to one of the Primitivo or Croatian clones than they are to other Californian clones. It would be unwise to assume a single migratory path radiating from an ancestral mother plant ought to be applicable to the clones.

Despite this ambiguity, the examination of SNPs, INDELs, transposable elements and other structural variants all support the derivation of all but one of the clonal selections from a common ancestral Zinfandel mother plant and show the accumulation of somatic mutations over time (Figs. 5 and 6). The structure of the Zinfandel genome, location of mutations among clones, their frequency and prevalence, and the relationship between these factors provides some insight into the nature of mutations in clonally propagated plants. Mutations among clones were predominantly heterozygous (Fig. 4) and uncommon heterozygous mutations shared by a subset of or individual clones were increasingly deleterious when they occurred in exons (Fig. 5e).

There are costs and benefits associated with clonal propagation [16]. Among the benefits are that the plants need not breed true-to-type; clonal propagation generally fixes heterozygous loci and valuable phenotypes. However, the increase in the proportion of deleterious alleles supports Muller’s ratchet, which posits that sex is advantageous and that clonal propagation increases mutational load [38]. Though these and previous data do not tell which mutations are actually recessive or dominant, they could remain hidden if they are recessive or do not manifest their deleterious effects [2,62]. However, even after taking into consideration the total length of exons, introns, and intergenic space (repetitive and non-repetitive), heterozygous mutations occurred at varying frequency in these regions and were least abundant in coding regions. The rarity of mutations in exons and commonality of mutations in repetitive intergenic space may have at least two components.

Mutations are likely more frequent in repetitive intergenic space as a result of the regulation of transposition by DNA methylation. Repetitive intergenic space had the highest rate of relatively unique SNPs and the ratio of transitions to transversions was significantly higher there than in other regions. DNA methylation is an important epigenetic control and is one mechanism that maintains genome stability and impairs the transposition of mobile elements [29,63,64]. Methylated cytosines, however, spontaneously deaminate faster than unmethylated cytosines [24,30]. Together, the expectations that intergenic regions are rich in transposable elements, that these regions are typically highly methylated and as a result will experience greater transition rates account for the high rates of SNPs in repetitive intergenic spaces among Zinfandel clones. Also notable, these data show that some transposable elements are not entirely silenced, with a substantial number inserting in genes or in close proximity to genes (Fig. 6c). These insertions could be effectively inconsequential or not; transposable element insertions can result in novel transcripts and affect gene expression regulation [11,65]. Gene body methylation is appreciated as a mutagenic “double-edged sword” [66], with benefits coming at the price. Recent work observed region-specific methylation in vegetatively propagated Sardinian white poplar that may serve an advantageous function [67] and others have suggested that the epigenome contributes to the success of vegetatively propagated plants [68]. Future work might also consider the long-term price associated with intergenic mutagenesis and the potential loss of methylation in vegetatively propagated plants.

The rarity of exonic mutations was surprising. After accounting for the length of these spaces in the genome and their repetitiveness, we expected uniform rates of mutation in exons, introns, and intergenic space. Instead, we observed that although rare somatic mutations in exons were increasingly deleterious, they were relatively scarce. Some degree of negative selection against deleterious variants in coding regions could explain why mutations were less frequent in coding than noncoding regions of the genome. The possibility of diplontic, clonal selection or competition between cell lineages that could purge otherwise consequential deleterious mutations has been modeled, but evidence of its occurrence is sparse [16,34,39]. The structures of apical meristems [35,69] and the tendency of somatic mutations to be heterozygous and recessive [2] place constraints on the likelihood that deleterious mutations would be subjected to negative selection. Periclinal divisions across cell layers could enhance diplontic selection [34] against dominant and/or hemizygous recessive alleles. Four and one half of Zinfandel’s genome is hemizygous; structural variations identified within the Zinfandel genome and the rampant hemizygosity reported in Chardonnay [10] could also expose otherwise hidden somatic variations to selective pressure hostile to the accumulation of deleterious mutations. Additional work should explore to what degree each of these factors, or others not considered here, explain why somatic mutations in exons were relatively infrequent and characterize the realized long-term consequences of mutation accumulation versus selection for grapevine and other clonally propagated plants.

## Conclusions

This study described the nature of the mutations causing the diversification of 15 clonally propagated grapevines and confirm their derivation from a single ancestral mother Zinfandel. The findings indicate that repetitive intergenic space, likely because of its higher rates of methylation in plants, is a significant contributor to the pool of mutations differentially observed among the clones. In addition, the analyses revealed that though relatively infrequent compared to intergenic mutations, mutations in exons were increasingly deleterious the less common they were among Zinfandel clones. This result is consistent with the expectation that vegetative propagation is associated with the accrual of mutations and adds that negative selection may simultaneously purge mutations from the genome. These findings add novel insight and nuance to our understanding of the nature and fates of mutations during vegetative propagation.

## Methods

### Zinfandel plant material and additional accessions

Fifteen Zinfandel clones were used for this study. Plants were confirmed to be clones of Zinfandel using the following microsatellite markers: VVMD5, VVMD7, VVMD27, VVMD31, VVMD32, VVMS2, VRZAG62, and VRZAG79 [44,70,71]. Fourteen of these clones are available through Foundation Plant Services (FPS) at the University of California Davis. Nine of the fifteen clones belong to the Zinfandel Heritage Vineyard Project, a collection of rare Zinfandel vine cuttings grown in the same vineyard. The identification numbers, common names, and source of the clones used in this study are listed in Table 1. An FPS identification number suffix of “.1” indicates that the clone underwent microshoot tip tissue culture therapy, with two exceptions. Pribidrag 13 and Pribidrag 15 are directly derived from the same plants as Pribidrag 4 and Pribidrag 5, respectively, but did not undergo microshoot tip tissue culture therapy. They are labeled with identical FPS numbers to make clear that the relationship between them is known. In this manuscript, Zinfandel clones will be referred to by the clone numbers and common names listed in Table 1.

### DNA extraction, library preparation, and sequencing

High quality genomic DNA was isolated from grape leaves using the method described in Chin *et al.* (2016) [53]. DNA purity was evaluated with a Nanodrop 2000 spectrophotometer (Thermo Scientific, Hanover Park, IL), quantity with a Qubit 2.0 Fluorometer (Life Technologies, Carlsbad, CA) and integrity by electrophoresis. For SMRT sequencing, SMRTbell libraries for the Zinfandel reference FPS clone 03 (Zin03) were prepared as described by Chin *et al.* (2016). For Illumina sequencing, DNA sequencing libraries for each of the fifteen Zinfandel clones were prepared using the Kapa LTP library prep kit (Kapa Biosystems) as described by Jones *et al.*, (2014) [72]. Final libraries were evaluated for quantity and quality using a Bioanalyzer 2100 (Agilent Technologies, CA). Zin03 SMRTbell libraries were sequenced on a PacBio RS II and Illumina libraries were sequenced in 100 and 150 base-pair paired-end reads on an Illumina HiSeq3000 sequencer (DNA Technology Core Facility, University of California, Davis). Genome sequences of additional *V. vinifera* were used in this study, including long reads from Cabernet sauvignon (NCBI BioProject PRJNA316730) and short reads from Cabernet franc, Chardonnay, Merlot, Pinot Noir, and Sauvignon blanc (NCBI BioProject PRJNA527006).

### Zinfandel genome assembly and annotation

*De novo* assembly of Zinfandel (Zin03) was performed at DNAnexus (Mountain View, CA, USA) using PacBio RS II data and the FALCON-unzip (v. 1.7.7) pipeline [53]. FALCON-unzip was used for its ability to assemble contiguous, phased diploid genomes with better resolved heterozygosity [53,73]. Repetitive sequences were masked prior to error correction using TANmask and REPmask modules in Damasker [74]. After error-correction (13,073 bp length cut-off), a total of 1.68 million error-corrected reads (N50 15Kbp, 29-fold coverage of expected genome size) were obtained and repeats were masked before overlap detection in the FALCON pipeline (v. 1.7.7). PacBio reads were assembled after testing multiple parameters to produce the least fragmented assembly. These conditions are listed in Additional file 4. Haplotype reconstruction was performed with default parameters. Finally, contigs were polished with Quiver (Pacific Biosciences, bundled with FALCON-unzip v. 1.7.7). Repeats were annotated on the Zin03 assembly using RepeatMasker (v. open-4.0.6) [75] and a *V. vinifera* repeat library [76].

The publicly available RNAseq datasets listed in Additional file 4 were used as transcriptional evidence for gene prediction. Each RNAseq sample was trimmed with Trimmomatic (v. 0.36; Additional file 4) and assembled with Stringtie (v. 1.3.3) [77] to reconstruct variety-specific transcripts. A detailed list of all experimental data used for the annotation procedure is in Additional file 4. This data was then mapped on the genome using Exonerate (v. 2.2.0, transcripts and proteins) [78] and PASA (v. 2.1.0, transcripts) [79]. Alignments, and *ab initio* predictions generated with SNAP (v. 2006-07-28) [80], Augustus [81], and GeneMark-ES [82] were used as input for EVidenceModeler (v. 1.1.1) [83]. EVidenceModeler was used to identify consensus gene structures using the weight reported in Additional file 4. Functional annotation was performed using the RefSeq plant protein database (ftp://ftp.ncbi.nlm.nih.gov/refseq, retrieved January 17th, 2017) and InteProScan (v. 5) as previously described [76].

### Genetic variant calling

Comparisons between Zinfandel clones and between Zin03 and other cultivars were made using the Zin03 genome as reference. This pipeline is described in Additional file 5. Small insertions and deletions (INDELs), single nucleotide polymorphisms (SNPs), and structural variations (SVs) were analyzed. The short Illumina reads belonging to the fifteen Zinfandel clones and additional cultivars were trimmed using Trimmomatic (v. 0.36; Additional file 4). Quality filtered and trimmed paired-end reads were then randomly down-sampled to 84 million (∼14X coverage) in each library to mitigate the possibility of sequencing depth-dependent outcomes. All libraries were aligned to Zin03 using bwa (v. 0.7.10) and the -M parameter [84]. For all genotypes, the median number of reads mapping to the Zinfandel reference genome was 97%. Next, Picard Tools (v. 2.12.1) were used to mark optical duplicates, build BAM indices, and validate SAM files (http://broadinstitute.github.io/picard). Variants were called using GATK’s HaplotypeCaller (v. 3.5) [85]. Then, called variants were filtered and annotated (--filterExpression “QD < 2.0 ∥ FS > 60.0 ∥ MQ < 40.0 ∥ MQRankSum < −12.5 ∥ ReadPosRankSum < −8.0”). Variant call files were combined using GATK’s GenotypeGVCFs. Having mapped Illumina reads corresponding to the Zinfandel reference onto itself, erroneous non-reference Zin03 calls (8.1%) were removed. The variants called included SNPs and INDELs.

Next, large structural variations among clones, between Zin03 and other cultivars, and between Zin03’s haplotypes were studied. First, Zin03 genes were compared to PN40024 and Cabernet Sauvignon (CS08) by mapping coding sequences on genome assemblies using Gmap (v. 2015-09-29) and the following parameters: -K 20,000 -B 4 -f 2. Hits with at least 80% identity and reciprocal coverage are reported. Genes annotated on Zin03’s haplotig assembly were also mapped to Zin03’s primary assembly to assess differences in gene content between Zin03’s haplotypes. SMRT reads from Zin03 and CS08 were mapped to Zin03 using NGMLR (v. 0.2.7) and structural differences were called with Sniffles (v.1.0.8) [55]. Zinfandel clones were compared to one another using Illumina short reads and Delly (v. 0.7.8) with default parameters [86]. The structural variations identified by Sniffles and Delly in Zin03 were intersected. Several filters were applied to the results of SV analyses. Transversions, non-reference Zin03 genotype calls, SVs annotated at the ends of contigs, and SVs that intersected the repeat annotation were filtered from Delly output.

### Transposon insertion analysis

PoPoolationTE2 (v. 1.10.04) [87] was used to identify transposon insertions in the Zinfandel clones; it was used following the workflow outlined in its software manual (https://sourceforge.net/p/popoolation-te2/wiki/Manual/). Insertions were called relative to Zin03 genome assembly and PN20024 [54]. As described in Kofler *et al.* (2016), PoPoolationTE2 analyses transposable element insertions and can identify novel and annotated TE insertions provided insertions fall within predefined families of TEs. The annotation produced by RepeatMasker was used for the analysis. In this manuscript, the TE insertions among the clones are reported using the classification system and nomenclature described by Wicker *et al.* (2007) [88]. In instances where the TE order and/or superfamily was not annotated, only the TE class and order, when available, are named in the associated figures and text.

### Relationships between Zinfandel clones

The relationships between Zinfandel clones were visualized by Principal Component Analysis and their relatedness was quantified (VCFtools v. 0.1.15) based on the method described by Manichaikul *et al.* (2010) [56]. This approach gives information about the relationship of any pair of individuals (unrelated, 3^rd^ degree relative, 2^nd^ degree relative, full siblings, and self) by estimating their kinship coefficient, which ranges from zero (no relationship) to 0.50 (self). These analyses used SNPs outside of repetitive regions.

## List of abbreviations

ZAP: Zinfandel Advocates and Producers
UC: University of California
FPS: Foundation Plant Services
SMRT: Single Molecule Real-Time
Zin03: Zinfandel 03
CS08: Cabernet Sauvignon 08
PCA: Principal component analysis
TEI: Transposable Element Insertions
SV: Structural variant
INDEL: Insertion/Deletion
SNP: Single Nucleotide Polymorphism

### Funding

This work was partially supported by start-up funds from the College of Agricultural and Environmental Sciences (UC Davis) to DC, the Louis P. Martini Endowment in Viticulture to DC and the NSF PGRP grant #1741627 to DC, MAW, and BG.

### Author contributions

AMV, MAW, BG, and DC designed the experiments. BBU, YZ, MMA, and RFB collected the biological material and generated the data. MAP carried out the chemical analysis of the clones. AMV, AM, YZ, DS, DL, and LKE analyzed the data. AMV and DC prepared the figures and wrote the manuscript. All authors contributed to the final version of the manuscript. All authors read and approved the final manuscript.

## Acknowledgements

We are grateful for the vision of the late James A. Wolpert, who established the original Zinfandel clone trials with the support of the Zinfandel Advocates and Producers (ZAP).

## Additional files

**Additional file 1.** .docx ; Method to extraction phenolic metabolites from Heritage Vineyard Zinfandel clones and discriminant analysis of Zinfandel clones based on their phenolic profiles.

**Additional file 2.** .xlsx ; Unique genes identified in Zinfandel, not identified in Pinot Noir and Cabernet Sauvignon (309), with associated Gene Ontology categories.

**Additional file 3.** .xlsx ; The first tab of this excel file is a summary of variants relative to the Zinfandel reference genome and the second is a summary of the SnpEff analysis of variants, with mean values ± SEM shown, and excluding sites where samples and Zin03 have identical heterozygous genotypes at the locus.

**Additional file 4.** .txt ; Settings and data used for Zin03 genome assembly, annotation, and variant calling.

**Additional file 5.** .sh ; Bioinformatic pipeline for SNP, INDEL, and SV calling.

